# Predicting the Impact of cis-Regulatory Variation on Alternative Polyadenylation

**DOI:** 10.1101/300061

**Authors:** Nicholas Bogard, Johannes Linder, Alexander B. Rosenberg, Georg Seelig

**Affiliations:** Department of Electrical Engineering, University of Washington; Paul G. Allen School of Computer Science, University of Washington

## Abstract

Alternative polyadenylation (APA) is a major driver of transcriptome diversity in human cells. Here, we use deep learning to predict APA from DNA sequence alone. We trained our model (APARENT, APA REgression NeT) on isoform expression data from over three million APA reporters, built by inserting random sequence into twelve distinct 3’UTR contexts. Predictions are highly accurate across both synthetic and genomic contexts; when tasked with inferring APA in human 3’UTRs, APARENT outperforms models trained exclusively on endogenous data. Visualizing features learned across all network layers reveals that APARENT recognizes sequence motifs known to recruit APA regulators, discovers previously unknown sequence determinants of cleavage site selection, and integrates these features into a comprehensive, interpretable cis-regulatory code. Finally, we use APARENT to quantify the impact of genetic variants on APA. Our approach detects pathogenic variants in a wide range of disease contexts, expanding our understanding of the genetic origins of disease.

---

Alternative polyadenylation is a ubiquitous regulatory process by which multiple RNA isoforms with distinct 3’-ends, and consequently differential stability, subcellular localization and translational efficiency, can be derived from a single gene (Figure 1A) (Di Giammartino et al., 2011; Elkon et al., 2013; Tian and Manley, 2017). APA is tightly regulated through a combination of cis-regulatory sequences -- most importantly a set of competing polyadenylation signals (PAS) -- and trans-acting RNA-binding proteins (RBPs) that recognize these sequences. Genetic variants that interfere with proper regulation of APA in cis have been implicated in disease (Danckwardt et al., 2008). For example, a mutation of a PAS in the *FOXP3* gene can cause IPEX, a rare disorder of the immune system (Bennett et al., 2001). Similarly, deletions or insertions in a PAS of the *Cyclin D1* gene have been associated with reduced survival of patients suffering from Mantle Cell Lymphoma (Wiestner et al., 2007).

The human genome supports on the order of 9 billion single-nucleotide variants and a comparable number of insertions and deletions; it is extremely likely that many more deleterious variants will be found that act by disrupting APA in cis. Experimentally characterizing the impact of every possible variant on regulation of APA is impossible given the low throughput of traditional experimental methods (Starita et al., 2017). Recently developed high-throughput measurement techniques such as massively parallel reporter assays (MPRA) can be used to quantify the functional impact of variants on gene expression (Findlay et al., 2014; Gray et al., 2018; Patwardhan et al., 2009), but even MPRAs are limited in throughput to tens of thousands of variants and are mostly targeted to a narrow subset of genes. Statistical methods such as genome-wide association studies have some success in linking genetic variation to disease but are limited to common variants. Overcoming these limitations of scale and generality requires accurate computational models that can be used to screen variants at genome scale and select for further investigation those variants predicted to be most disruptive.

Recent work in computational biology, often using deep learning approaches, has taken steps towards building predictive models of regulatory processes. Examples include DeepBind (Alipanahi et al., 2015), Basset (Kelley et al., 2016), DeepSea (Zhou and Troyanskaya, 2015), and even a recent model of APA (Leung et al., 2017). However, the quality of such functional models is determined not only by the underlying algorithm and model architecture but by the quality and size of the training data. The use of very large libraries of synthetic reporter constructs with targeted variation can overcome intrinsic size limitations of biological datasets and have proven to be highly effective in training models that are sensitive to the complexities of gene regulation (Rosenberg et al., 2015). However, such approaches have not yet been applied in the context of APA, a process that, in spite of its importance, remains relatively understudied.

To generate a dataset large and diverse enough to represent the complexity of the APA code we constructed and transiently expressed minigene libraries of more than 3 million unique UTRs and obtained the isoform and cleavage data from the expressed RNA (Figure 1B). In total, we assayed over 250 million degenerate bases, 3.6-fold more than are contained in all 3’UTRs from the human reference genome. Our synthetic random PASs are expressed within a smaller library of host UTR sequences, allowing us to measure, compare and learn from a diverse set of contexts. We trained a Deep Neural Network (DNN), APARENT, on the random 3’UTR library in order to predict alternative isoform expression of each library member given its sequence. We show that APARENT accurately predicts isoform expression on entirely held-out libraries. We adapted and applied various DNN visualization techniques to identify important features of the APA regulatory code, including sequence and secondary structure motifs, their position-dependence, and higher-order interactions between them. By fine-tuning the network with a small number of endogenous examples, APARENT was able to predict alternative isoform expression in native human genes to a far greater extent than a DNN trained on endogenous data alone. Finally, we demonstrate that APARENT can identify deleterious mutations that disrupt gene function by mis-regulating APA.

## Constructing a Library of over 3 Million APA Reporter Constructs

Most APA occurs between PASs expressed in the 3’UTR; the simplest and most common form being 3’UTRs with only two APA signals (Elkon et al., 2013). A PAS is defined by the 6-base central sequence element (CSE), most commonly AATAAA, and its upstream and downstream sequences (USE, DSE). To survey the impact of cis-regulatory elements on APA, we constructed 12 reporter libraries with 3’UTRs variable in length, location and extent of randomization, PAS-to-PAS distance, and sequence context (Figure 1C). Contexts used for library construction include 3’UTR sequences from seven human genes each known to natively express two 3’UTR PASs with canonical or near-canonical CSEs, including a membrane translocase (*TOMM5*), one tRNA synthetase (*AARS*), a kinase (*ATR*), a heatshock protein (*HSPE1*), a long non-coding RNA (*SNHG6*), a transcription factor (*SOX13*), and one pseudogene (*WHAMMP2*).

The libraries derived from the *TOMM5* 3’UTR express a relatively weak proximal PAS (pPAS) (CSE = TATAAA) in competition with the stronger, canonical AATAAA at a distal site approximately 300 bases downstream. We separately randomized two adjacent sites upstream of the pPAS then included either wild type sequence or an additional 20-bases of randomization downstream, resulting in a total of four distinct libraries derived from the native *TOMM5* 3’UTR sequence. In the six shorter UTRs, we randomized 25 bases each in the USE and DSE, centered on the cleavage site (Figure 1C).

**Figure 1.**
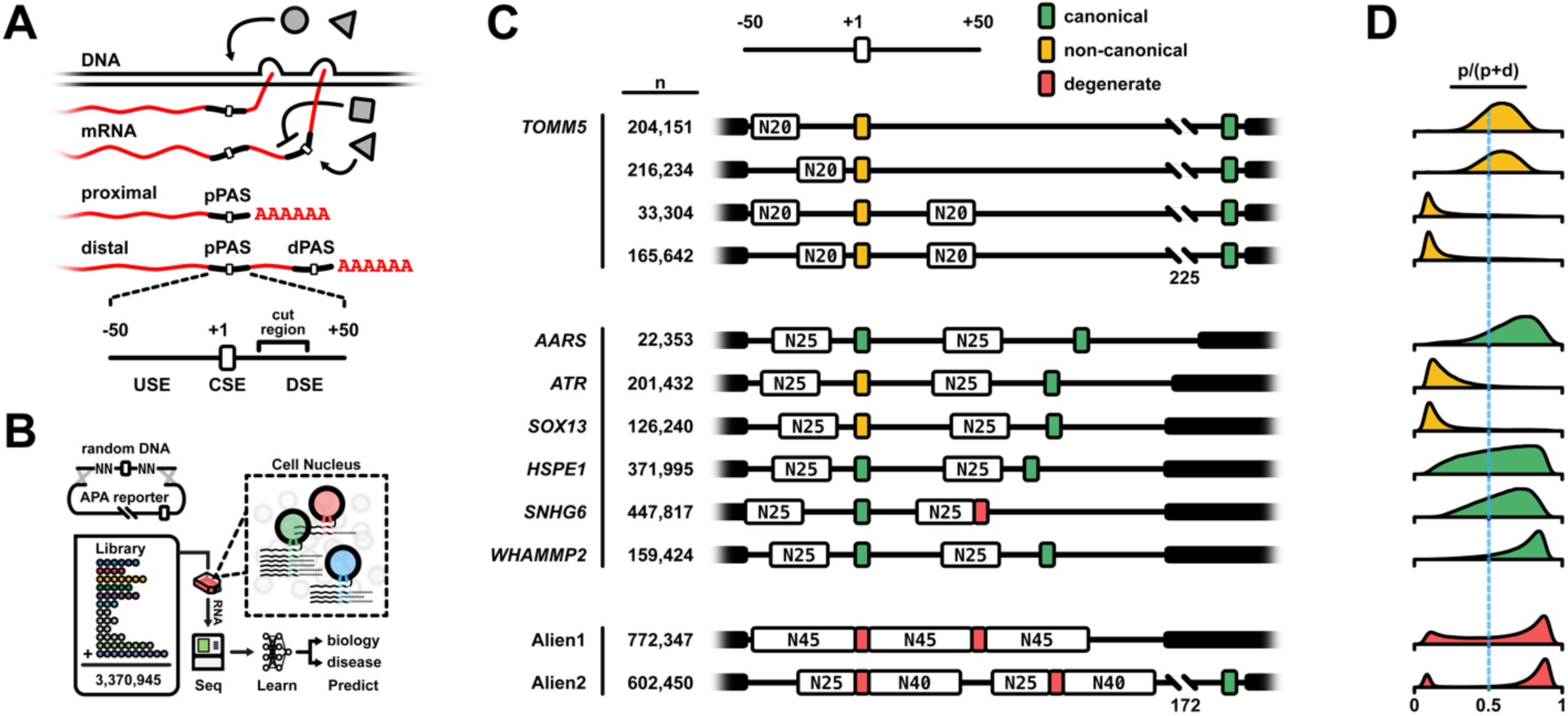
Massive Parallel Reporter Assay for Alternative Polyadenylation. (A) Newly-transcribed mRNA is targeted by multiple factors (grey) that enhance or suppress selection of alternative polyadenylation sites. A PAS (below) is defined by the 6-base Central Sequence Element and regions of approximately 50 bp both upstream and downstream. (B) Millions of unique reporters are cloned from degenerate oligos and transiently transfected in human cell culture where they are expressed and alternatively polyadenylated. RNA is extracted, sequenced and quantified for every reporter. These data are then used to train, validate, or test the model. (C) The library is comprised of multiple sublibraries that vary in structure and 3’UTR context (human italicized; CSE sequence denoted in legend). Degenerate sequence was introduced up and/or downstream (N20-45) of the proximal PAS, and in some cases, within the CSE (degenerate, red). The total number (n) of unique members for all libraries is over 3 million. (D) The distribution of relative proximal site usage per unique member for each library. Each histogram is color-coded to match the proximal CSE. See also Figure S1.

Additionally, we constructed fully alien 3’UTR libraries by inserting doped CSEs (degeneration biased for functionality) within fully randomized sequence. The goal of these non-native libraries was to understand to what extent a putative APA code could be generalized to, or even learned from, entirely synthetic sequence contexts. Alien1 expresses two consensus AWTAAA CSEs -- each having an equal probability of being the stronger AATAAA or the weaker ATTAAA -- in close proximity. All sequence 45 bases upstream, between, and downstream is fully randomized. Alien2 expresses a 120-base degenerate 3’UTR with two A-rich alternative hexamers (95% A, 2% G/C, 3% T) expanding the variety of non-canonical CSEs.

## A Massively Parallel Reporter Assay for Alternative Polyadenylation

To quantify isoform expression, we transiently transfected the library in HEK293 cells, extracted the expressed library RNA, and sequenced it (Figure 1B). In total, we collected isoform expression data for 3,370,945 unique reporters with an average of 38 reads per reporter. The majority of our reads, 53%, mapped to proximal isoforms while 28% mapped to distal isoforms. Native expression from the genes used for the human-derived libraries have a similar proximal bias (58%; Müller et al., 2014). The remaining reads (19%) originated from de novo polyadenylation signals within the degenerate regions.

Isoform ratios exhibited a wide range of variation within and between libraries (Figure 1D, S1). Variation could be attributed to the CSE but also the surrounding regulatory sequences. In UTRs with two signals - both AATAAA - the proximal site was slightly preferred, consistent with the “first come, first served” hypothesis of kinetic coupling (Bentley, 2014; DeZazzo and Imperiale, 1989) (Figure 1D; *AARS, HSPE1, WHAMMP2*). A weaker CSE can retain preference, especially if the distance to the alternative site is relatively large, as with the *TOMM5* UTR. Notably, randomization of the *TOMM5* DSE results in nearly complete suppression of the proximal isoform. The native sequence we randomized is T-rich (40% T), a conserved feature of DSEs (Nunes et al., 2010; Pérez Cañadillas and Varani, 2003). *TOMM5* library members filtered for random DSEs enriched for T restored proximal expression (.58 relative pPAS usage for T >= 40%, .06 for T < 40%), suggesting that the feature does indeed promote site selection, and underscoring the importance of regulatory sequence outside of the CSE.

## Predicting Alternative Polyadenylation with APARENT

Next, we turned to training a deep neural network model for predicting PAS usage from DNA sequence. To construct our model we assumed that competing APA sites are regulated independently, a reasonable assumption if the pPAS and dPAS are sufficiently separated such that the proximal site DSE does not overlap with the USE of the distal site. Consequently, to learn a general model of APA regulation, the model needs to be trained only on randomized proximal sites (and non-randomized distal sites).

APARENT (Figure 2A), was trained to predict the ratio of proximally to non-proximally polyadenylated isoforms of each variant UTR given its sequence as input. We trained APARENT on data from 9 out of 12 synthetic 3’ UTR libraries. More specifically, 95% of the data from these 9 libraries was used for training (~2.4M variants), 2% for validation (~50,000 variants) and 3% for testing (~80,000 variants evenly sampled from each library). Three randomly selected libraries were left out entirely, allowing for testing of the final model in a sequence context unrelated to any context seen during training.

**Figure 2.**
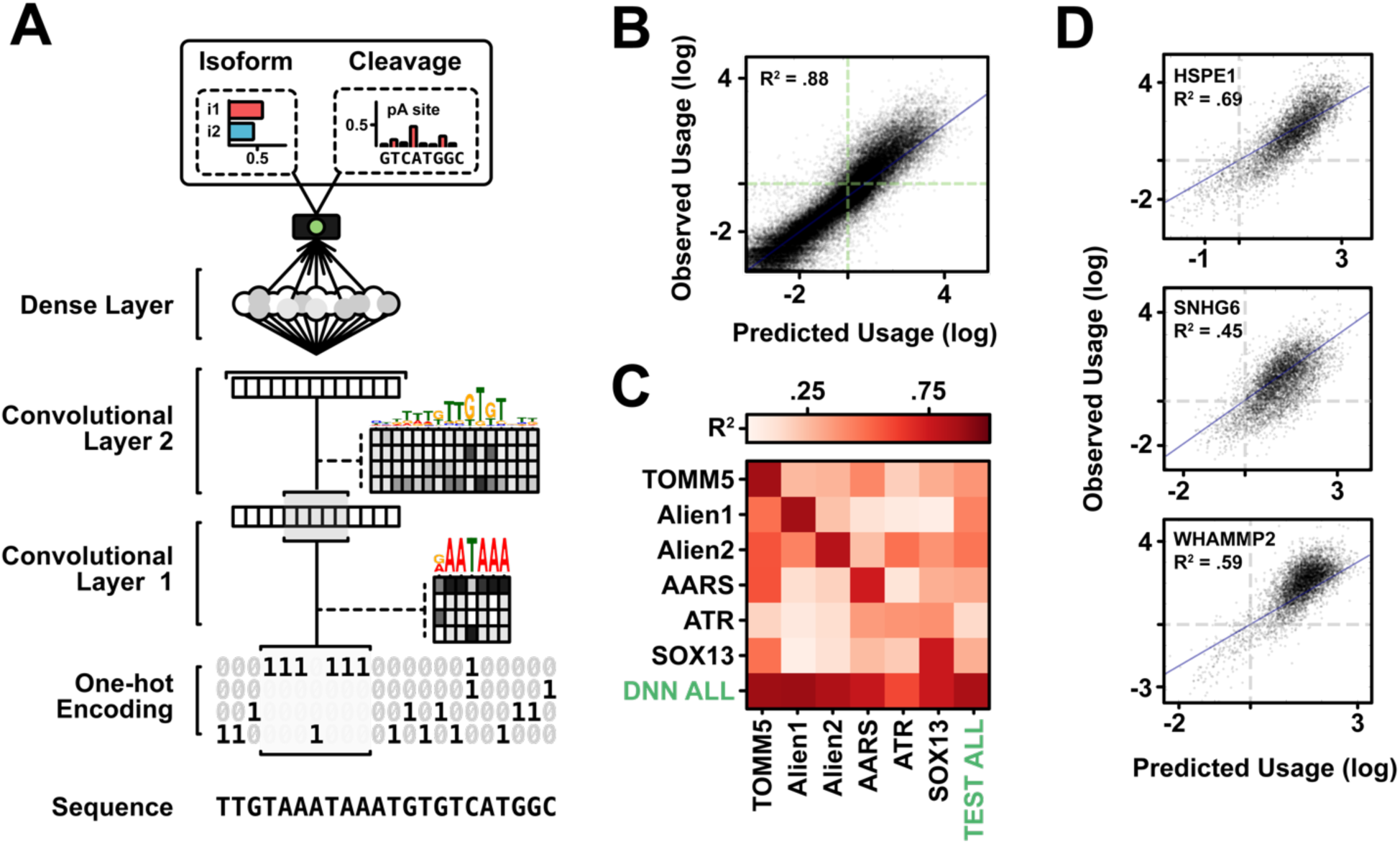
Model Architecture and Performance. (A) An illustration of APARENT, a CNN consisting of two convolutional layers, a maxpooling layer (not shown), one fully-connected (dense) layer, and a modular output layer. The input sequence, which is a UTR library variant aligned on the proximal CSE, is passed as a one-hot-encoded two-dimensional matrix to the model. APARENT can generate two possible outputs: % proximal isoform abundance and % cleavage at each nucleotide position. (B) Scatter plot of predicted vs. observed proximal isoform logodds (the log of the odds of % proximal isoform selection), evaluated on the held-out test set (~80,000 sequences) of the trained-on UTR libraries using APARENT (R^2^ = 0.88). (C) Cross-library confusion matrix. R^2^ is measured on predicted vs. observed proximal isoform logodds. The diagonal entries represent tests where the training set and test set come from the same plasmid library, while off-diagonal entries are tests of models trained on one library but tested on data from a different library. (D) Scatter plots of predicted versus observed proximal isoform logodds on held-out plasmid libraries not trained on by the model (HSPE1, SNHG6, and WHAMMP2; mean R^2^ = 0.58). See also Figure S2.

The best-performing model consisted of two convolutional layers interlaced with subsampling layers, a fully connected layer, and a logistic regression output node on top. To evaluate the model, we tested its ability to infer proximal isoform selection (as a result of pPAS usage). When tasked with predicting proximal isoform logodds (the log of the odds of proximal isoform selection), the DNN performed remarkably well on the combined test set (R^2^ = 0.88; Figure 2B). Furthermore, to validate that the best APA model is obtained from joint training across all libraries, separate DNNs were trained on each individual library and cross-tested against the test sets from every other library (Figure 2C). R^2^ is measured between predicted and observed proximal isoform logodds. The DNN trained on the combined data (bottom row) performed at least as well, and in some cases better, than every individual network did on its corresponding library. As a baseline test, we also compared the DNN model to a 6-mer linear logistic regression model trained on the combined library and found that the DNN significantly outperforms this simpler model, suggesting that positional as well as non-linear effects are important for accurately predicting APA (Figure S2A).

Finally, we asked whether the model could generalize to new UTR-specific contexts by predicting the proximal isoform logodds on the three held-out libraries (*HSPE1*, *SNHG6* and *WHAMMP2*). The predictions correlate well with the observed measurements (mean R^2^ = 0.58, Figure 2D, S2B), suggesting that the regulation learned by the DNN is not limited to the gene contexts used during training.

## Sequence Determinants of Isoform Selection

Deep neural networks have in recent years been applied extensively to sequence-based genetic data and different methods have been proposed for visualizing and quantifying sequence features learned by these networks. By building on such techniques from computational biology and combining them with approaches from computer vision, we developed a way of visualizing the sequence determinants identified by the network to affect PAS preference, not only in the first convolutional layer but also in higher layers, as well as ranking them by order of importance.

First, convolutional filters of the first layer were visualized as position weight matrices (PWMs) (Figure 3A, S3A-B). The position-specific effect on PAS usage of each filter was quantified by measuring the correlation between activations and isoform usage at each position across the PAS (Cuperus et al., 2017). Negative correlation corresponds to a negative impact of the motif on site preference, while positive correlation suggests an enhancing effect. Next, we cross-referenced the PWMs with published binding data, as well as the Compendium of RNA Binding Protein motifs (Ray et al., 2013) using the Tomtom comparison tool (Gupta et al., 2007), and surveyed the top-scoring results for APA mediators (Figure 3A).

**Figure 3.**
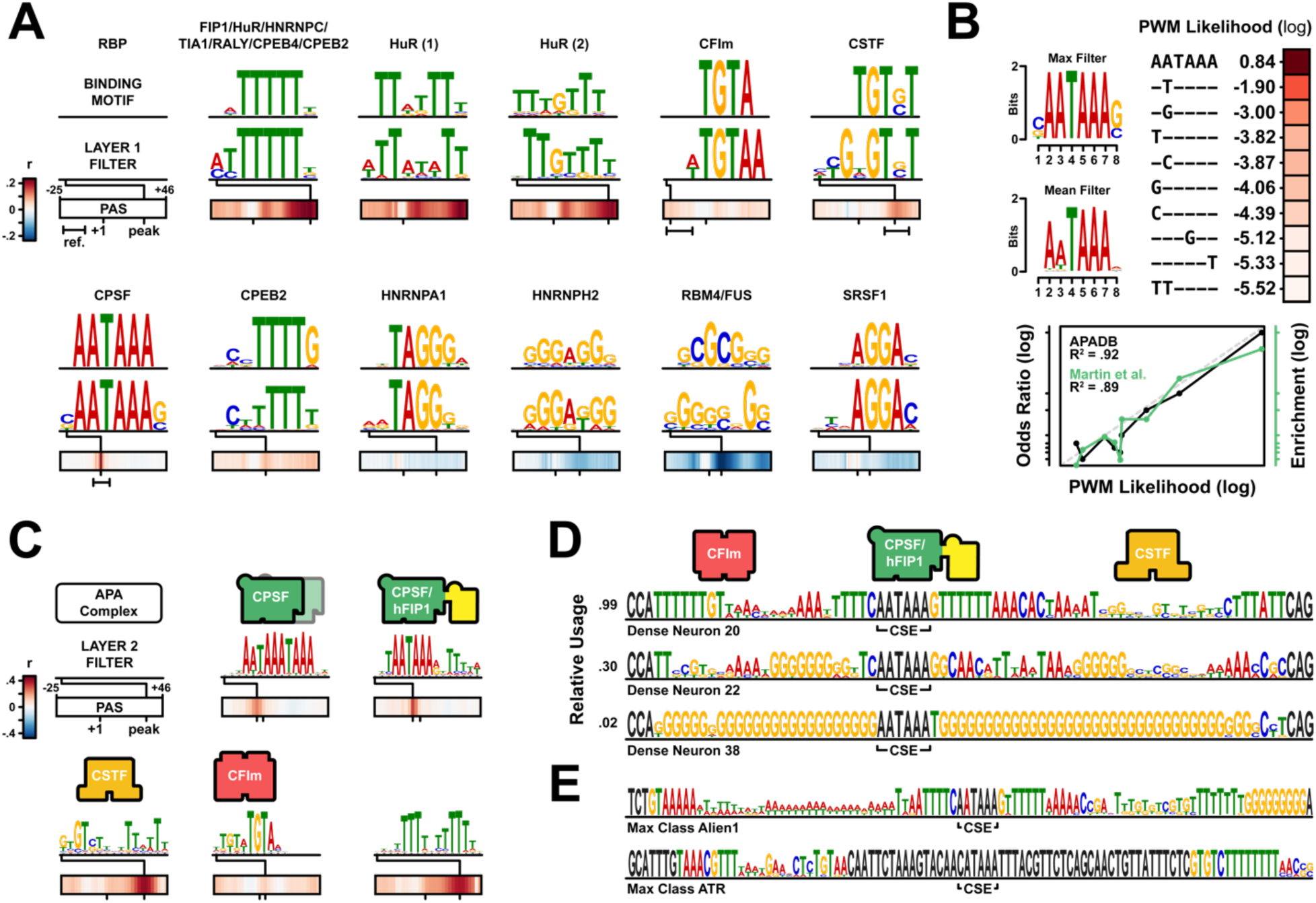
Layer-by-Layer Feature Analysis Parallels Increasing Complexity of the PAS. (A) RBP binding motif logos and positional effects for eleven select layer 1 convolutional filters. Positional effect is measured by Pearson’s coefficient of correlation between filter activation and proximal isoform logodds. The start position of peak correlation is annotated as a line passing through the heatmap. Published positional data (Martin et al., 2012) are marked as range bars below the heatmap. The canonical CSE, CSTF, and CFIm motifs and their positional effects are replicated in an independent random model initialization (Figure S3B). (B) Scoring CSE Strength. Left: Sequence logos for the maximally-activated CSE detector filter and its mean-activated filter when sampling 40k random input sequences and scaling their PWM contribution by the filter activation magnitude. Right: Ranked CSE variants extracted from the PWM. All but one are single mismatches from the canonical AATAAA. Below: Correlation between APARENT’s scoring and previously-published experimental data (solid lines trace the published rank order). (C) Visualization and positional effects for five layer 2 convolutional filters. Proposed effector interactions are illustrated above each filter logo. (D) Sequence logos of individual dense neurons from the fully-connected layer optimized for maximal neuron activation (black = non-variable sequence). The dense neuron most associated with high site usage prediction resembles a complete PAS involving interactions with several components of the polyadenylation machinery. (E) Optimization for proximal site selection in two library contexts. See also Figure S3, Table S1-2, and Movie S1.

We identified matches for all components of the core polyadenylation machinery that directly interact with cis-regulatory sequences (i.e., CFIm25, CPSF160 and Wdr33, and CstF64) (Schönemann et al., 2014). Importantly, these filters are activated maximally when the motif occurs at the expected position in the PAS. Moreover, the enhancing or suppressing effect assigned to each filter is consistent with the known regulatory role of its matched RBP. For example, filters clearly resembling the canonical CSE, AATAAA, are found multiple times and their activations highly correlate with increased isoform use when positioned in a narrowly-defined region at the center of the PAS. TGTA, a motif known to recruit the APA enhancer CFIm, is also found in multiple filters that are positively correlated with proximal upregulation, in particular when located in the USE (Gruber et al., 2012; Zhu et al., 2018). GT-rich motifs in the DSE have a strong positive correlation with pPAS selection, consistent with recognition by CSTF (Pérez Cañadillas and Varani, 2003; Takagaki and Manley, 1997). The polyT motif is found to be strongly correlated with proximal selection at two distinct locations in the DSE. Immediately downstream of the CSE, the enhancing effect and position agrees well with the binding site of Fip1 (Kaufmann et al., 2004; Martin et al., 2012). At further distances from the CSE, the enhancing effect could be attributed to multiple factors with known sequence preference to polyT, such as HuR, RALY, TIA1, CPEB2/4, and HNRNPC (Nunes et al., 2010; Pérez Cañadillas and Varani, 2003; Tian and Manley, 2017). Many additional filters, potentially shaped by interactions with known APA effectors, were also identified (Figure 3A, HNRNPH2, SRSF1, RBM4, etc.). For example, HNRNPH2 has been proposed to suppress APA site selection by physically blocking CSTF from binding (Nazim et al., 2017). This mechanism is supported by our model, which identifies the HNRNPH2 binding site as a repressor mainly in the DSE. Filters that did not match to binding sites for known RBPs, possibly suggesting novel regulatory interactions, were also found (Table S1).

The motifs identified by the DNN were validated directly in the data (without the use of any model) using a logodds ratio analysis, where every possible 6-mer occurring in the USE, CSE, and DSE were scored and ranked according to the expected increase in odds ratio of proximal isoform selection (Rosenberg et al., 2015). The top scoring enhancer-and repressor sequences found (Figure S2C) agree well with the most influential motifs extracted from the convolutional filters of the DNN.

## Quantifying the Strength of the CSE

The CSE regulates APA by directly binding CPSF160, CPSF30, and Wdr33 (Clerici et al., 2017, 2018; Sun et al., 2018). Variants of the consensus hexamer AATAAA, typically differing by a single base, are functional and common (Gruber et al., 2016). These variants exhibit reduced affinity for recruiting the CPSF complex, with the degree of affinity determined by the nucleotide substitution. Given that the CSE is randomized in some of the UTR libraries, the DNN should be able to detect these functional variants and give appropriately scaled responses to reflect CSE strength. To test this, we generated new convolutional filter PWMs where each contributing sequence was scaled by the magnitude of the filter response (Figure 3B, top). The scaled PWM indeed captures the relative strength relationship between variants, as the order and magnitude of the top 10 PAS variants extracted from the PWM are nearly identical with APADB values and a separate independent study (Gruber et al., 2016) (Figure 3B, bottom).

## Visualizing Motif Interactions

Next, we constructed consensus sequence logos for the motifs learned in the second convolutional layer, as well as measured their positional effect on isoform selection. Since layer 2 filters cover more of the input sequence than layer 1 filters, the resulting APA code decomposition consists of longer sequence motifs. In general, the second convolutional layer captures longer repetitions of short motifs, such as T- or G-stretches wider than the filters of the first layer, or combinations of distinct elements (Figure 3C, S3C-D, Table S2). The composition and positional effect size of these combinations aligns well with the current understanding of APA regulation. For example, the DNN identifies the core hexamer AATAAA in combination with a downstream polyT motif to substantially upregulate site selection compared to the average effect of the core hexamer AATAAA, reflecting known observed interactions between Fip1 and other components of the polyadenylation machinery (Kaufmann et al., 2004). The DNN also identifies strong combinatorial effects of DSE determinants, such as GT-rich motifs (CSTF-binding) preceding a polyT motif further downstream. One filter is sensitive to dual TGTA motifs, supporting the notion that CFI binds as a dimer (Yang et al., 2010, 2011). Finally, one of the showcased filters appears to be sensitive to a “multi-PAS” consisting of two overlapping canonical hexamers forming the 10-mer AAWAAAWAAA, estimated by the model to increase proximal site selection compared to a single AATAAA.

## Global Determinants of Isoform Selection

Next, we turned to the problem of understanding the global relationship between USE and DSE regions. A common approach in image classification networks for visualizing higher-layer DNN features is to apply gradient-based optimization to find input images that maximally stimulate neurons in various layers. For example, (Simonyan et al., 2013) optimizes input images that maximize the final class probability score. Here we develop a similar optimization algorithm suitable for sequence-based neural networks that finds sequence patterns which maximally activate a particular neuron in the network (Figure S3E).

The optimization routine is first carried out on the dense layer (the fully-connected layer preceding the final output layer). Starting from 50 random UTRs from the Alien2 library, the sequences were optimized with respect to maximal activation of a neuron in the layer. The optimized sequences, when stacked in a sequence logo, create a visual representation of the global structure that the neuron is sensitive to (Figure 3D, S3F-G). The neurons clearly specialize on different sequence subspaces, where G- and GC-rich sites are much less favored compared to sites containing various conserved T- and GT-rich stretches. By inspecting the bottom two logos, inclusion of the polyG motif in mainly the DSE, but also in the USE, is an efficient mechanism for suppressing a site. Similarly, from inspecting the top logo, having polyT-stretches at the ends of the USE and DSE regions appear to be associated with strong proximal selection. Also, a polyA motif ending with a single C approximately 5-10 bp downstream of the CSE is present in all of the high-proximal neurons.

We also analyzed sequences optimized for maximum proximal expression. Starting from random input encoded with a particular UTR library background sequence, we generated 50 optimized sequences per library and stacked them into sequence logos. The resulting logos revealed global sequence elements associated with strong proximal selection (Figure 3E, S3H-I). A time-lapse animation of an optimization (Movie S1) exemplifies how the procedure populates the PAS with stronger enhancers first and settles on some motifs only after the other motifs have already emerged, suggesting combinatorial effects. The USE and DSE regions exhibit very similar characteristics across libraries, suggesting there is a “consensus” template for a strong PAS. Strong upstream regions typically include runs of polyT separated by a non-conserved A-rich sequence, and possibly a TATA, polyC, or TGTA motif. Strong downstream regions typically include a stretch of polyT, an A-rich stretch, followed by GT- or CT-rich content and finally another stretch of polyT. The GT-/CT-enriched stretch is located 20-30 bp downstream of the CSE, and the trailing polyT sometimes contains a single A. While the generated patterns share many similarities, there are subtle differences due to the library template bias. An example is the polyG track found in the 3’ end of the Alien1 DSE. Immediately downstream is a competing polyA site encoded in the template. The network appears to have learned to find an optimal equilibrium point where it fills the 5’-end of the DSE with enhancing motifs, thus promoting usage of the proximal site, while filling the 3’-end with polyG (a verified suppressor), repressing the nearby distal USE. In the *ATR* context, expressing a fixed non-canonical CSE, optimization settles on multiple TGTA motifs in the USE. The techniques developed here thus make it possible to generate a global view of how multiple sequence elements interact with one another to determine isoform choice.

## Predicting Cleavage Distribution with APARENT

RNA-seq not only makes it possible to distinguish between proximal and distal isoforms but even provides information about the precise cut position. We thus asked whether our model could be trained to predict the probability for cleavage occurring at any given position. To directly learn the cleavage distribution of each variant we used a DNN almost identical to the isoform-based model. The main difference is that instead of predicting a ratio between two isoforms, the final layer of this network outputs a multinomial probability distribution of cleavage at any position across the entire sequence (Figure 4A). The output cleavage distribution can easily be translated to an isoform ratio; the probabilities of the positions included in the polyA isoform of interest are added and normalized by the sum of all positional probabilities. The network was trained using identical training-, validation- and test-set splits as used earlier.

**Figure 4.**
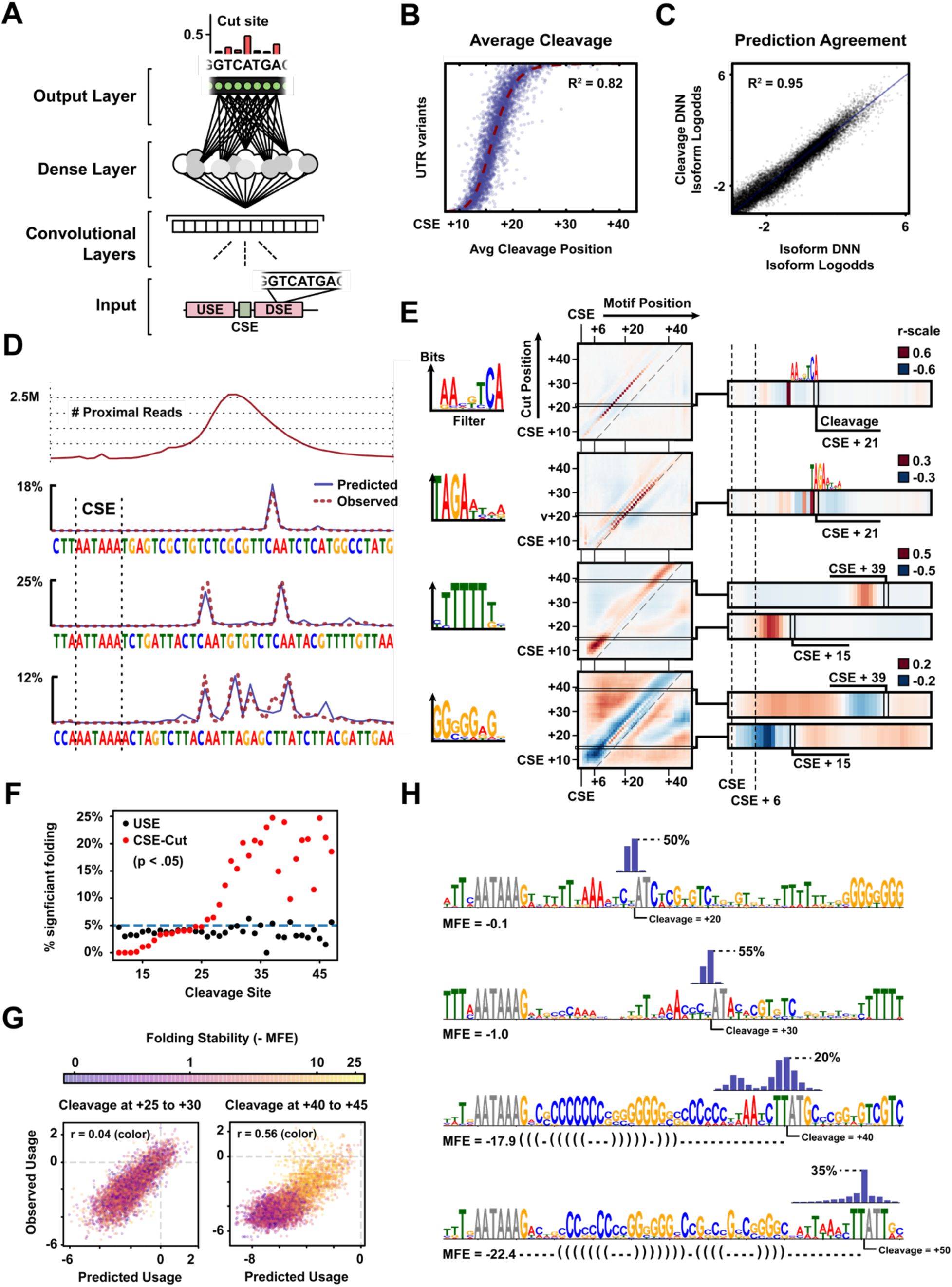
Cutsite Prediction and Analysis. (A) APARENT’s architecture for predicting cutsite distribution. The output module is replaced with a 186-way multinomial probability vector (obtained from a Softmax activation layer), where each probability node corresponds to the % Cleavage occurring at the nucleotide of the same position in the input sequence. Cut probabilities across an arbitrary DSE are illustrated. (B) Average cleavage position prediction results on the test set of the Alien1 UTR library. The X-axis denotes the average cleavage position of a particular variant, and the Y-axis lists each test set variant, sorted on observed average position. (C) Correlation of predictions between the isoform-based output layer and integration of cleavage distribution across the proximal cut region. (D) Three example sequences from the Alien1 test set, overlaid with the predicted cleavage distribution (blue) and observed cleavage distribution (red). The panel also includes a plot of the total number of polyadenylated reads of the Alien1 library, aligned vertically on the example sequence positions. (E) A selection of filters validating known (CA dinucleotide, top) and newly discovered determinants guiding cleavage. Each plot measures correlation between a filter activating at a certain position and cleavage occurring at some other position. The first two filters display cleavage site dinucleotides identified by the network. To correctly interpret the cleavage site position in the plot, one must account for the in-filter position of the motif. For example, the high-intensity heat of the CA-filter occurs 6 nucleotides upstream a cleavage site, merely due to the fact that CA resides at an offset of +6 inside the filter. The remaining two filters display impactful cleavage site regulators, namely polyT/polyG motifs. Figure S4B displays additional identified motifs, including the presumed binding site of CSTF. (F) The relationship between secondary structure and cleavage distance. Sequences upstream (black) and downstream (red) of the CSE were folded in silico and the significance of their secondary structure evaluated as the fraction against 100 control sequences (mono-and dinucleotide shuffling of each input) having lower MFE. The relative amount of members where the fraction of sequences with stable folding was greater than 95% of the shuffled variants was then plotted against average cleavage position. (G) Predicted and observed site usage of Alien1 sequences at cleavage positions relatively near (left) or far from (right) the start of the CSE. The color encodes MFE of the DSE regions. Predicted (and observed) usage correlates with MFE only at the far region of cleavage. (H) Sequence optimization for predefined cleavage positions. APARENT was initialized with random sequence input (except for a hard-coded CSE and cleavage site) and iteratively optimized for cleavage at four increasingly distant positions (20, 30, 40 and 50 nt downstream of the CSE start position). Sequence intervening the CSE and cleavage site were folded. Shown are the optimized sequence logos, the average cleavage distribution (bar plot), the average MFE, and the predicted structure (dot-parent notation) for 20 optimized sequences at each cleavage position. See also Figure S4 and Movie S2.

We evaluated this generalized version of APARENT in two ways. First, we compared the predicted average cleavage position with the observed average position of every test set variant (Figure 4B). The two quantities correlated well (R^2^ = 0.82 for Alien1, R^2^ = 0.55 for the held-out *WHAMMP2* library). Second, we compared the area under the proximal region of the predicted cleavage curve, which corresponds to total isoform abundance, against the previously predicted proximal isoform ratios (Figure 4C). The two predictions show strong agreement (R^2^ = 0.95), meaning that the predicted distributions identify cleavage sites of previously unseen (test set) UTRs while preserving the relative magnitude of isoform expression.

Encouraged by the substantial cleavage variation observed in the libraries, we searched for UTR variants with extreme deviation in cleavage sites. These UTRs reveal a rich landscape of possible cleavage distributions ranging from unimodal cuts, to bimodal distributions separated by as much as 10 nucleotides, to a cluster of positions where cleavage is initiated at many different sites (Figure 4D). Remarkably, APARENT is able to predict these sites with high precision, suggesting the choice of cleavage site itself is governed by a deterministic regulatory code.

## The Determinants of Cleavage Site Selection

To identify the sequence determinants and positional effects influencing the choice of the cleavage site, we applied APARENT to a random sample of 120,000 sequences from the Alien1 library, predicting their entire cleavage distribution. Each Layer 1 filter activation at every position was recorded and correlated with the magnitude of cleavage at every position, resulting in set of two-dimensional plots that together with their consensus sequence logos describe the regulatory impact of each filter motif on every cleavage site as a function of position (Figure 4E, S4A-B).

Consistent with earlier studies, the DNN filter heatmap identifies the CA dinucleotide as the most favored substrate for cleavage (Derti et al., 2012). However, the TA and GA variants are also identified as functional, albeit weaker cleavage sites, supporting experimental evidence and conservation analyses suggesting that CA is not essential (Chen et al., 1995; Li and Du, 2013). To validate these findings, we extracted every dinucleotide in the region +5 to +35 bps downstream of the PAS from the entire UTR library and recorded the percent cleavage occurring at each dinucleotide. Using these measurements, we calculated the logodds ratio of cleavage at each dinucleotide (Figure S4C). The CA, TA, and GA dinucleotides showed a considerable increase in cleavage likelihood compared to other elements, with CA at the top of the ranking.

The GT-rich motif, typically identified as the CSTF binding site (Pérez Cañadillas and Varani, 2003), is found to enhance cleavage when located immediately downstream of the cleavage site (Figure S4B, Movie S2). Interestingly, the heatmap indicates that the effect of the GT-rich motif is only functional for cleavage sites located ~15-30 bp downstream of the first nucleotide of the CSE. Finally, the DNN characterizes the polyG and polyT motifs as highly effective cleavage site regulators; polyG immediately preceding a cleavage site suppresses its usage, while an upstream polyT motif enhances usage. The correlation coefficient of cleavage for the polyT motif is almost as strong as the dinucleotide CA. Even more interesting, according to the heatmaps, polyG and polyT enhance or suppress cleavage at sites either adjacent to or far away from the CSE, locations at which the dinucleotides CA, TA, and GA have no measurable impact.

Recently, 3’ mRNA secondary structure was implicated in guiding cleavage site position (Wu and Bartel, 2017). Curious whether the feature was present in our data, we computationally folded the upstream and downstream sequences of over 100k Alien1 pPASs and evaluated the significance of structure relative to each cut position (Freyhult et al., 2005). We found that cleavage position, beyond an optimal distance downstream of the CSE, does indeed correlate with increased secondary structure (Figure 4F). We then probed whether APARENT had learned this property. First, we evaluated the relationship between cleavage position and DSE minimum free energy (MFE). For cleavage sites near the CSE, no correlation was found (Figure 4G). However, for cleavage at distances >40 bases, APARENT’s predictions were highly correlated with MFE, in agreement with the results of Wu and Bartel.

Next, we used APARENT to optimize randomly initialized UTR sequences for cleavage at various distances from the CSE (Figure 4H, S4D-F, Movie S2). For cleavage sites within 30 bases, the optimized DSEs converged to sequences with no predictable structure. Rather, the model positioned core processing elements (CSTF, various RBP binding sites, etc.) in optimal locations relative to the cleavage site. These optimal motif locations agree with the heat intensities of the filter analysis (Figure 4E). For example, the presumed binding site of CSTF is placed immediately downstream of the cleavage site, and for very short cleavage sites, PolyG is inserted at the far-3’ end of the DSE. Beyond this region of cleavage, APARENT instead favored sequences upstream of the cleavage site that are predicted to form stable hairpins with stem lengths that increase monotonically with site distance.

## APARENT Accurately Predicts Native APA

Next, we turned to APADB, a public dataset of APA events from multiple human tissues. APA events found in native genomic context contain a larger degree of complexity than what is captured by the libraries used for training our models. Specifically, in the libraries the sequence of the dPAS is mostly fixed and in relatively close proximity to the pPAS. In a native context, both distal site composition and distance varies greatly. However, we hypothesized that individual PAS strengths are independent when signals are well-separated, and that site distance adds only a prior towards proximal site selection. If we further assume that the same sequence-specific regulation governs both proximal and distal sites, the DNN should be able to score both of them. Hence, the task is reduced to learning a linear function that maps the predicted score of each PAS, and their log-distance, to the isoform ratio. This linear function is learned by optimizing for the logodds of the measured APADB isoform ratios. Since our model predicts the ratio of one isoform versus another, pairs of adjacent APA sites within UTRs of the APADB dataset were considered. We further filtered for a sufficiently high read count and removed sites where the CSE differed by more than two bases from the canonical hexamer (filtered n = 674). The final model predicts a relative ratio between APA isoforms A and B even when the entire regulatory context, including the distal PAS, is varied (Figure 5A). We used leave-one-out cross-validation to give an unbiased estimate of the model performance.

**Figure 5.**
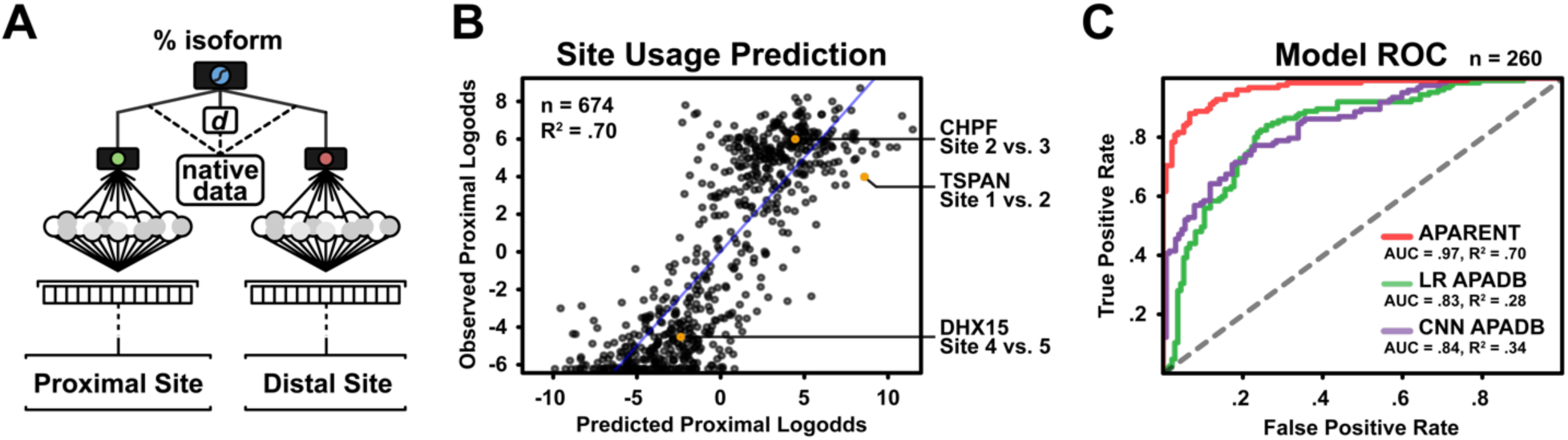
Performance of APARENT on Endogenous APA. (A) An illustration of the extended model architecture when predicting the relative isoform ratio between adjacent APA sites of the APADB dataset. The network, from the first convolutional layer up to the final output regression node, is replicated and placed over both the proximal- and distal APA sites. The predicted isoform logodds score of each network is used as input, together with the log of the site distance, to a regression node that is optimized to predict the target isoform ratios of APADB. (B) Correlation between the isoform logodds predictions of the extended model and the observed values from the APADB dataset. Leave-one-out Cross-validation is performed on the entire filtered dataset (n = 674). (C) ROC curves obtained on the classification task of predicting whether the proximal isoform or distal isoform of an APA event is selected for the most. Predictions were performed on a held-out APADB test set (n = 260). APARENT’s classification performance is compared against an identical network trained exclusively on the APADB data. A baseline 6-mer regression model is also tested. See also Figure S5.

The predicted isoform logodds were strongly correlated with the observed values for held-out APA events (R^2^ = 0.70, Figure 5B). Performance also correlated with APADB read coverage, indicating that the prediction performance increases monotonically with higher-quality estimates of the true isoform ratios (Figure S5A). When reducing the problem to predicting whether an event is mostly proximally or distally polyadenylated, APARENT is shown to significantly outperform an identical DNN trained exclusively on the endogenous APADB data, with an AUROC (area under the receiver operating characteristic curve) of 0.97 (Figure 5C, S5B). These results indicate that the millions of synthetic APA events contain more information about the regulation of APA than what can be extrapolated from naturally occurring events alone.

## APARENT Predicts SNVs Linked to APA Misregulation

To assess the DNN’s ability to predict the impact of variants on APA in the human genome, publicly accessible RNA-Seq data from the GEUVADIS project (Lappalainen et al., 2013) were used together with the 1000 Genomes reference sequences to obtain a set of variants occurring near polyadenylation sites. We first identified relevant genes containing APA events using the Tandem 3’UTR annotation compiled by the MISO project (Katz et al., 2010). Next we searched the 1000 Genomes data for human samples with single-nucleotide variants (SNVs) occurring in the 3’UTR within 75 bp of an annotated PAS. The variant sequences were recorded and the corresponding RNA-Seq data was processed using MISO to estimate true isoform ratios of every APA event. Since the entire UTR sequence, including the distal site, is identical between wild type and variant samples except for the SNV, the shift in isoform logodds predicted by the DNN should be directly comparable to the observed shift in isoform logodds estimated by MISO from the RNA-Seq data. The logodds predictions correlate highly with the estimates (R^2^ = 0.68, Figure 6A).

**Figure 6.**
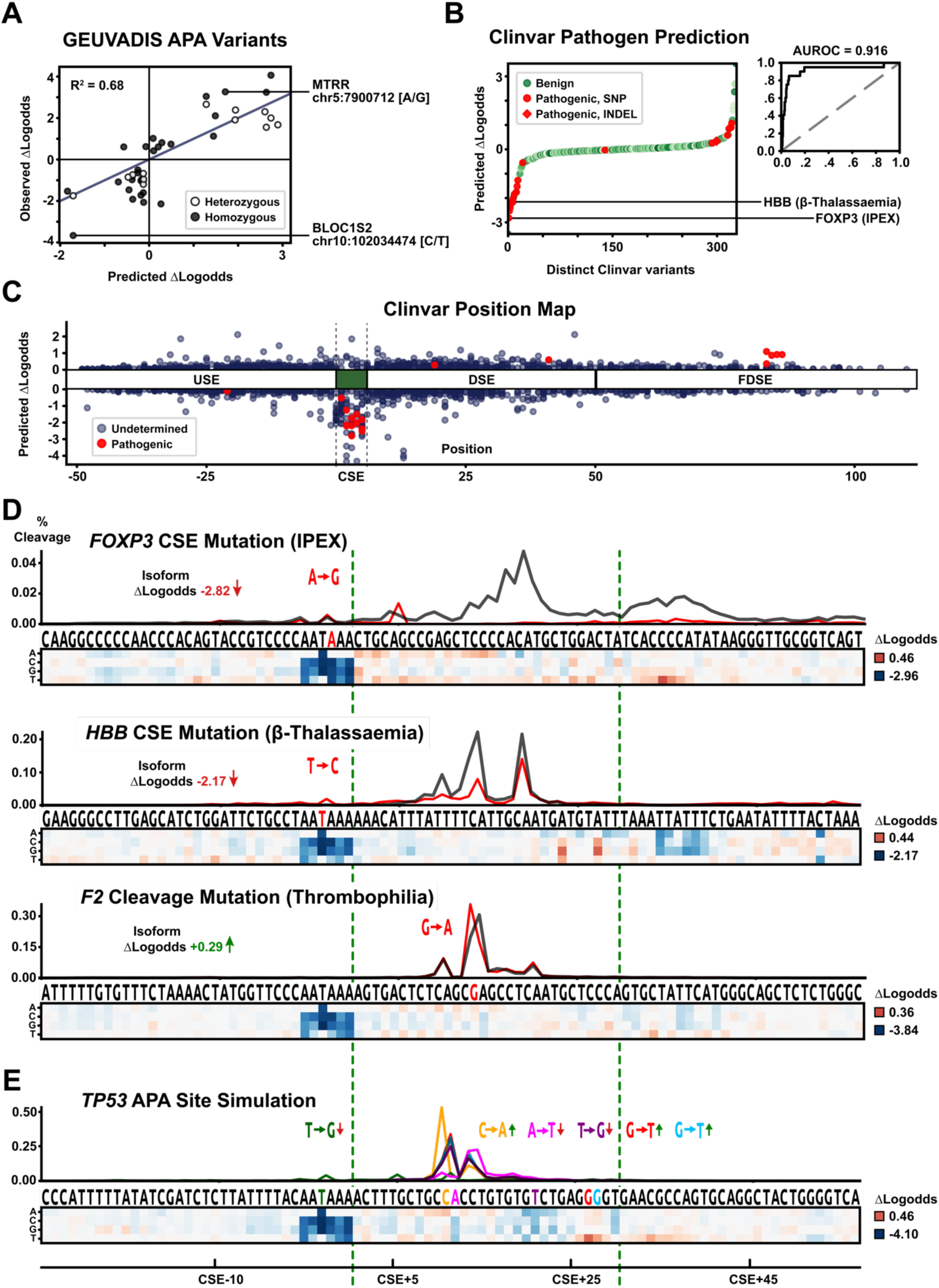
Predicting SNV Effects. (A)Correlation of SNV impact between predictions (change in isoform logodds) and GEUVADIS measurements. Variants mapped to one strong and one weak PAS are indicated. (B) Classification performance on ClinVar mutations (SNVs and InDels) when using APARENT’s predicted change in isoform logodds as a proxy for pathogenicity. Two disease-associated SNPs are indicated. (C) Scatter plot of predicted change in isoform logodds vs. position relative to the CSE. Pathogenic (red) and unknown (blue) variants are annotated (FDSE = Further Downstream Sequence region). (D) Visualization of the predicted cleavage distribution for pathogenic SNVs in human 3’UTRs. Predictions for the wildtype (black) and mutant (red) distributions are shown with the nucleotide substitution above. The effect on isoform expression (cleavage occurring between the green dashed lines) is indicated below the gene name. A heatmap depicts the effect of each potential SNV by coloring each nucleotide-position entry according to the predicted change in isoform logodds (range shown at right). To see the effect of USE and DSE SNVs more clearly, the color intensities of the CSE hexamer mutations have been downscaled by a factor of 2. (E) Simulation of six potential SNVs in the disease-associated *TP53* 3’UTR. The cleavage distributions are displayed over the sequence (color matched to the variant bases). Effect sizes for every potential SNV on isoform selection are shown in the heatmap. The color intensities of the CSE hexamer mutations have been downscaled by a factor of 2. See also Figure S6.

Confident in our model, we investigated whether mutations occurring in human PASs could be linked to disease. We mapped ClinVar data -- SNVs and InDels linked to medical phenotypes (Landrum et al., 2018) -- to PASs in APADB and tasked APARENT with predicting changes in isoform expression due to each variant. The majority of variants annotated as pathogenic were predicted to have substantial isoform shifts (Figure 6B). More than half of the pathogenic variants map to the CSE, which APARENT considers to be especially sensitive to variation (Figure 6C). APARENT also predicts that a small cluster of pathogenic DSE variants cause shifts in isoform expression, suggesting that non-CSE mutations cause misregulation. However, a considerable amount of mutations remain to be classified; many predicted to have strong effects on isoform expression. Given the high classification performance (AUROC = 0.916), the model could prove a useful tool for filtering unclassified mutations for candidate variants with high likelihood of causing disease by affecting APA.

Two specific SNVs have previously been identified as causal for disease by disrupting the CSE of the pPAS (Figure 6D, top). The first mutation transforms the canonical CSE AATAAA of the *FOXP3* UTR into a much weaker variant, AATGAA, which substantially lowers the cleavage magnitude at the downstream CA dinucleotide (predicted isoform logodds change = −2.82). This mutation has been linked to IPEX syndrome (Bennett et al., 2001). In our analysis, we also simulated every possible mutation across the PAS, illustrated as a heatmap below the sequence. In the *FOXP3* UTR, the DNN finds that any disruption to the dual-CA dinucleotides in the DSE, serving as cleavage sites, strongly represses isoform expression. Conversely, enriching the sequence further downstream with T enhances expression.

The second mutation, which occurs in the pPAS of the *HBB* 3’UTR and has been linked to beta-thalassaemia, disrupts the CSE AATAAA by transforming it into the variant AACAAA (Orkin et al., 1985). All other SNVs at this CSE linked to beta-thalassaemia are also predicted to alter isoform expression (Giordano et al., 2005; Jacquette et al., 2004; Jankovic et al., 1990; Rund et al., 1992; Waye et al., 2001). In the DSE, G/T substitutions upregulate selection significantly, possibly by creating CSTF binding sites (Figure 6D, middle). Additionally, an important polyT motif is found further downstream and substitution for any nucleotide other than T within this motif represses usage. The DNN also identifies a cryptic CSE ACTAAA in the DSE as a strong competitive signal. Given that beta-thalassaemia is caused by APA misregulation in the *HBB* UTR, our model suggests there are many more mutations beyond that of CSE variation which could misregulate *HBB* 3’-end processing. Interestingly though, in contrast to the *FOXP3* UTR, none of the cleavage sites are identified as downregulating selection if disrupted by an SNV. We suggest that the saturation of CA and TA dinucleotides in the DSE, in combination with a long polyT cleavage enhancer immediately succeeding the CSE, safeguards the PAS from misregulation due to a point-mutation on any single cleavage site.

We then challenged APARENT with predicting the impact of a variant (20210 G>A) at the cleavage site in the prothrombin gene (*F2*) 3’UTR. This gain-of-function variant is found in nearly 2% of the population and has been directly linked to the blood disorder thrombophilia (Danckwardt et al., 2008; Gehring et al., 2001). The single base substitution is directly adjacent to the cleavage site and converts a weak CG dinucleotide to CA resulting in a 1.45-fold increase in mRNA processed at this site in a reporter assay containing a competing PAS. APARENT accurately identifies the weak wildtype CG dinucleotide as a functional cleavage site, and furthermore, predicts that the CA variant increases use of the site about 1.3-fold (Figure 6D, bottom). However, APARENT also predicts that other variants including those in the CSE would result in a more pronounced shift in isoform distribution.

Finally, we assigned APARENT the task of predicting the impact of every possible SNV in a *TP53* PAS implicated in disease (Figure 6E, S6A-B) (Stacey et al., 2011). The heatmap identifies two important regulatory motifs in the DSE: the CSTF binding site TGT[C/G]T, whose disruption downregulates selection significantly, and the HNRNPH binding site, whose disruption upregulates selection. The heatmap also shows the importance of the unique CA dinucleotide, disruption of which shifts cleavage to a non-canonical cleavage site and overall lowers affinity. Also, notice that the DNN correctly weighs the relative effect of each mutation over the CSE, the least significant mutations being AATAAA>ATTAAA and AATAAA>AGTAAA, and the most significant being AATAAA>AAGAAA.

## DISCUSSION

We introduced a deep neural network model capable of accurately predicting isoform distribution for alternative polyadenylation in human 3’UTRs. Our model predicts both the usage of a particular PAS relative to all other PASs in a 3’UTR and the exact position of cleavage and polyadenylation with single-nucleotide resolution. Many of the sequence features identified as important by the model could be mapped to known cis-regulatory sequence elements. For example, our model learned sequence motifs consistent with known binding sites of core components of the polyadenylation and cleavage machinery (CFIm, CSTF, CPSF and Fip1) as well as other RBPs previously implicated in regulation of APA. Importantly, our model also identified regulatory features that have not previously been described. For example, we found that the choice of cleavage site is regulated deterministically by sequence features besides the favored CA dinucleotide. The DNN identified GA and TA as almost equally functional, but also found that cleavage site selection is regulated by a more extensive cis-regulatory code combining RBP binding motifs (e.g. polyG/polyT/CSTF) with secondary structure.

Our work is to our knowledge the first to build meaningful, biologically interpretable visualizations of higher-order features learned in deep layers of sequence-based neural networks. Recent work on visualizing the motifs learned by the first layer of a CNN has begun to address latent skepticism about the interpretability of such models compared to linear k-mer regression or k-mer SVM models. We expand on this work and show that our network identifies important regulatory features of widely varying complexity, ranging from short motifs learned in the first layer, to longer and spatially connected combinations of motifs in the second layer, to full-length PAS compositions, including secondary structure, learned in the deeper layers. Identifying determinants and non-linear interaction of that complexity would be impossible with a naive k-mer kernel, in particular with regard to long sequence motifs that require exponentially many k-mer weights. A non-linear model alleviates this problem of exploding dimensionality, and our visualizations overcome the lack of interpretability.

A practical application for APARENT comes from its ability to quantify the impact of genetic variants on APA isoform distribution. We validated our model using variants from the 1000 genomes project for which associated RNA-seq data was available. We then applied APARENT to all 3’UTR variants in ClinVar within 50 base pairs of the CSE and found that many variants annotated as pathogenic also had a strong impact on APA, suggesting a molecular mechanism for disease. We identified a large number of variants classified as variants of unknown significance that nonetheless resulted in strong shifts in the APA isoform distribution, making them intriguing candidates for future investigation. Furthermore, APARENT’s sensitivity in predicting the impact of disease-linked variation in less-characterized elements, like the cleavage dinucleotide, demonstrates the model’s understanding of a range of 3’ processing events.

The performance of deep neural networks and other machine learning approaches dramatically increases with the size of the available training data. In order to fully leverage the computational power of DNNs, models were trained on data from a massively parallel reporter assay consisting of over three million APA mini-gene reporter constructs with randomized sequences. In part, the choice of randomized libraries was one of convenience because very large numbers of constructs can be created in a single cloning step and at the cost of a single oligonucleotide. However, the randomized sequence elements are also free from any inherent biases that might exist between native elements or the genes that host them. Furthermore, the random sequences, with effects ranging from negative to neutral to positive, uniformly probe the phenotypic landscape which potentially benefits a model’s ability to identify impactful variations amongst a large background of silent mutations. Still, by targeting variation to a large number of native sequence contexts and to different regions relative to the core hexamer, we captured some of the regulatory context present in human 3’UTRs.

In future work, we intend to build on our results in several ways. First, though we focused exclusively on APA in the 3’UTR, PASs are also found in terminal introns. A more comprehensive model of APA would thus need to capture not only the cis-regulatory determinants of APA but also the interplay between APA and splicing. Although predictive models of alternative splicing are available, more work is required to integrate them with the models introduced here. A modified library, where variation is embedded near a terminal exon and spliceoform expression is assayed alongside APA, should refine our understanding of these interdependencies. Second, it is likely that some of the cis-regulatory sequences identified in our assays differentially control transcript stability rather than regulating APA directly. Such sequence elements could be identified in an independent MPRA focused on mRNA stability, either via updated libraries that integrate variability in regions differentially included in alternative isoforms, or in modified assays that incorporate expression over time (Rabani et al., 2017). Third, several genes exhibit differential site usage when expressed in different cells and tissues. For example, pluripotent stem cells typically prefer proximal sites, whereas terminally differentiated neuronal cell-types have a strong bias for distal sites and long 3’UTRs (Zhang et al., 2005). It will be interesting to perform similar APA MPRA measurements in other cell types in order to better understand cell-type specific contributions to APA. Finally, models like APARENT allow for preliminary analyses to identify potentially significant candidate variants. Genetic screens, first conducted in silico, could aid in the design of more informative experiments. Still, the work presented here provides a comprehensive view of the regulatory code guiding APA and has direct implications for both basic biology and genomic medicine.

## Acknowledgements

We thank J. Shendure, P. Sample, B. Groves, A. Carignano, A. Kuchina, S. Pochekailov, A. Baryshev, S. Chen, A. Khakhar, R. Lopez, S. Rao, C. Roco, B. Wang, Y. Zhang, E. Wilson, M. Hirano, R. Langevin, A. Lin, and N. Clarke for their valuable advice and discussions. This work was supported by NIH R01HG009136 to GS.

## Author Contributions

N.B., J.L., A.R., and G.S. conceived and developed the project. N.B. conducted biological experiments. N.B. and J.L. developed the models and conducted computational experiments. N.B., J.L., A.R., and G.S. performed analyses and wrote the paper.

## Methods

### Cloning

For the *TOMM5* library, a vector was constructed by replacing the bGH pA signal with an IDT gBlock designed with the *TOMM5* 3’ UTR sequence. The vectors were then linearized at the site of randomization and assembled with Klenow-extended degenerate oligos that overlapped at the pPAS site and included compatible overhangs. For all other libraries, two oligos were designed to anneal at the pPAS and included 25-45 bases of randomization and UTR-specific overhangs. Matching overhangs specific to each UTR were added to a linearized reporter by inverse PCR. All libraries were constructed using Gibson Assembly. Library sizes were estimated by plating a small amount of transformation and extrapolating based on colony counts. The remaining transformants were grown in 50mL overnight culture. Individual plasmid libraries, and individual clones from each library, were Sanger sequenced to confirm the expected structure and diversity.

### Cell Culture and RNA Extraction

HEK293 cells were grown on ECM-coated plates in DMEM supplemented with 10% FBS and antibiotics. Cells were trypsinized, washed with PBS, and lysed 36-48 hours post-transfection. mRNA was purified with the NEB polyA purification kit that utilizes T20-linked, magnetic beads.

### Sequencing Library Construction

Polyadenylated RNA was reverse-transcribed with an anchored polyT primer containing Illumina adapter sequences and a unique molecular identifier (Adapter-UMI-T18VN). Library cDNA was then amplified using a library-specific forward primer containing additional Illumina adapter sequences (Adapter-Library-FWD) and reverse primer matching the adapter sequence added during RT. Amplification was conducted and monitored with a qPCR instrument and stopped early to minimize PCR biases.

### Barcoding and Mapping

For libraries with PASs relatively close together, we sequenced and searched 5’ to 3’ across both signals for the site of polyadenylation. For longer UTRs, we identified the polyadenylation site via reverse-complement mapping of sequence adjacent to the paired-end poly-T read.

To record full library sequences, including those downstream of the proximal cleavage site, we sequenced amplicon prepped directly from the plasmid library. The first 20-25 degenerate bases upstream of the proximal PAS was used as a barcode to enable mapping of cleaved mRNA back to the full UTR sequence. Sequencing reads were long enough to precisely locate cut sites for all proximal and some distal isoforms. The resulting dictionary of sequences was used to map reporter RNA back to the originating plasmid sequence.

### Defining the Site of Polyadenylation

For all libraries, with the exception of TOMM5 and Alien2, both proximal and distal polyadenylation sites were identified with a single, sense-strand sequencing read that primed directly upstream of the randomized USE. We used the open-source adapter trimming software package cutadapt v1.15 (Martin 2011) with polyA trimming parameters (-a ‘A’*18-m 2). For TOMM5, exact polyadenylation positions were identified only for the proximal isoform while distal cleavage was inferred from reads that lacked polyadenylation within the proximal DSE. For Alien2, the sequence downstream of T18 in the anti-sense read was mapped back to full-length library members.

### Data Processing

Having collected sequencing reads from all MPRAs, the raw data was filtered, clustered and transformed into a set of well-defined 3’ UTR libraries. Full-length RNA reads were filtered on a sufficiently high quality, after which they were clustered on the randomized region upstream of the pPAS, resulting in a dictionary of sequence variants for each library. The dictionary was further expanded by sequencing the plasmid library to include members that expressed no distal isoform. RNA reads were mapped to each respective dictionary entry by matching the upstream region with shortest hamming distance.

For each mapped read, the Polyadenylation cleavage site was determined by scanning the read for the Poly-A tail. For some libraries, reads can map to a non-degenerate distal site found 2-300 bases downstream of the degenerate proximal site. The cleavage positions of all the mapped reads for one particular sequence variant was stored as a vector associated with that variant. Reads mapping to far-away (non-random) distal sites were recorded in a special “distal” position of each count vector. The resulting dataset consisted of a per-library dictionary of unique sequence variants with associated cleavage-position count vectors. The library datasets were passed through a final filtering step to select for high-confidence variants. This step removes sequences that are supported by fewer than 10-20 unique-UMI RNA reads (the exact number depends on the specific library and its read coverage), or sequences that contain >75% A-nucleotides in a 12-20bp region (to safeguard all libraries against internal priming artifacts).

In order to jointly model all variants of the combined multi-library, they need to be aligned in a meaningful way; the regions of randomization as well as the position of polyA signals vary for each library. Consequently, each variant was padded with wildtype sequence such that the most proximal CSE is located at equal relative positions across libraries. The resulting consolidated sequence template contains 50 nucleotides upstream and 130 nucleotides downstream of the pPAS and allows any learning algorithm to unambiguously relate the target response variable (“proximal usage”) to a specific sequence element (namely the CSE at position 50).

### Data quantification

The raw input data to the learning models consisted of the 186bp-long aligned library sequences, coupled with identifiers specifying origin library. Depending on the learning model used, this raw input data was featurized in different ways: For the neural networks, the input sequences were One-hot-encoded as two-dimensional matrices. For the logodds ratio analysis and K-mer regression, the sequences were transformed into long sparse vectors that encode which K-mers are present. These model-specific featurizations are covered in detail in subsequent sections.

The raw output (or target) response data revolves around the cleavage-position count vector C^v^ associated with each sequence variant v (obtained from the library processing described in the previous section). This vector records at each position (relative to the location of the proximal CSE) the number of RNA reads supporting cleavage-and polyadenylation (or isoform selection) at that position. From this vector, a number of different statistics were obtained and used for training and testing the models. These statistics are explained below:

#### Proximal Isoform Ratio

The fraction of RNA reads supporting the isoform obtained from usage of the proximal PAS CSE. Since cleavage typically occurs in a range (not just one position) downstream of the selected CSE, an aggregate of RNA read positions was used to determine proximal isoform support. In agreement with previous studies, we defined [PAS+10, PAS+35] as the range of cleavage positions supporting the proximal isoform (where PAS denotes the start position of the CSE hexamer), and all other positions support competing, non-proximal, isoforms. Given the 186-bp long cleavage count vector C^v^ and the count c^v^_dist_ of cleavage counts occurring at a far-away distal site, the proximal isoform ratio P^v^_Prox_ for sequence variant v is estimated as:

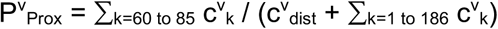

Due to the library alignment, this measure is unambiguous and relatable to the proximal PAS CSE at position 50 for all libraries, while nearby randomly inserted PASs will be considered competing signals, as their cleavage likely maps to locations outside the 60-to-85 proximal window.

#### Proximal Isoform Logodds

The log of the odds of proximal PAS usage (or equivalently, the log of the odds of selecting the proximal isoform). The log odds scale transforms the response variable from the closed range [0, 1] to the open infinite range (-inf, inf) and makes it more suitable for statistical analysis such as regression, correlation tests etc. The statistic is calculated as:

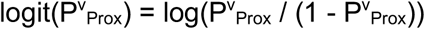

#### Cleavage Distribution

This multinomial response variable normalizes the vector of cleavage counts for each variant such that it sums to 1 and thus defines a probability distribution of cleavage occurring across the 186-bp long stretch in the UTR. The Cleavage Distribution P^v^ for variant v is calculated as:

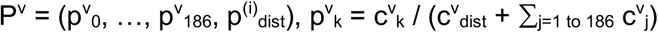

The cleavage distribution vector retains much more detailed information about cleavage-and polyadenylation for a particular variant than the proximal isoform ratio; the vector records with nucleotide resolution the probability of cleavage. This information is lost with the proximal usage ratio, which is computed by aggregating the probability of cleavage over multiple positions.

### Logodds Ratio

To validate the sequence motifs and regulatory effects on APA learned by the models, a statistic called the logodds ratio was estimated for every 6-mer located in the USE-and DSE regions (Figure S2A). The statistic was also estimated for every possible 6-mer occurring at the exact location of the CSE hexamer. For each K-mer S, the logodds ratio LOR(S) is calculated as:

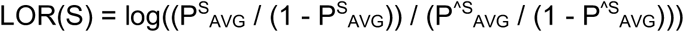

where P^S^_AVG_ is the average proximal isoform ratio of library variants containing S in the region of interest. P^^S^_AVG_ is the average ratio of variants not containing S. The logodds ratio is an estimation of the expected increase in the odds of selecting the proximal isoform over non-proximal isoforms given the presence of a K-mer. In Figure S2A, the natural log is used.

A similar analysis was performed to estimate the affinity for cleaving at each of the 16 dinucleotides (Figure S4C). For each sequence, the positions in the cleavage distribution vector P^v^ with non-zero cleavage probability were recorded along with the corresponding dinucleotides where the cleavage occurred. The average cleavage probability PC^S^_AVG_ was calculated from the subset of cleavage probabilities that were recorded with dinucleotide S. Similarly, PC^^S^_AVG_ was averaged from the remaining set of cleavage probabilities. These estimates were passed to the logodds ratio analysis LOR(S) as defined above. To estimate the certainty in the calculated values, 50-fold Bootstrapping with replacement was used to obtain a 95% confidence interval.

### Training and Test splits

For all learning models, 9 of 12 libraries were used as training data while the remaining 3 libraries were kept as independent test libraries. However, for each of the 9 training libraries, a small test set was kept. The exact training- and test splits are given below:

1. The 3 libraries HSPE1, SNHG6 and WHAMMP2 were held out entirely from training.
2. Each of the 9 remaining libraries were shuffled individually as follows:

a. Each individual library L was sorted in ascending order based on read count.
b. For each individual library L, every odd-numbered variant (in the sorted order) was bucketed into subset L_A_ and every even-numbered variant was bucketed into subset L_B_. The sort order was then altered by concatenating the entire subset L_B_ after L_A_. As an effect, half of the high read count-variants are placed in the lower half of the library sort order and the other half is placed in the upper half. This operation balances having high-read count variants in both training- and test sets.
3. The 9 libraries were shuffled together as follows:

a. Starting from the end of each library’s sort order (where the highest read count-variants are placed), the next variant to add into the back of the combined library is chosen in round-robin order from each individual library, until they run out of variants. As an effect, the back of the combined library (where the test set will be taken from) contains an equal number of sequences from each library, thus balancing the test set with regard to library representation.
b. The training set (95% of the combined library) is taken from the beginning of the library sort order. In total, the training set contained more than 2.4 million variants. The test set (3% of the combined library, taken from the very end of the library sort order) contained more than 80,000 variants. The remaining 2% of the data was used as a validation set and contained more than 50,000 variants. The validation set was used for cross-validation in K-mer regression and for Early stopping when training the neural networks (to prevent overfitting).

### Statistical tests

The Pearson correlation coefficient was used whenever we measure correlation between two variables and are interested in whether they are correlated or anti-correlated. The Pearson correlation coefficient r, for two sample sequences {xM_i_}_i = 1 to n_ and {y_i_}_i = 1 to n_, is calculated as:

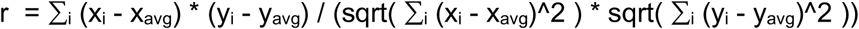

Throughout the paper, R^2 is used as a measure of correlation when comparing observed output response variables to values predicted by our models. R^2 is a common statistic to use for measuring goodness of fit for predictive models. Note that here R^2 is calculated as the squared Pearson correlation coefficient (r^2). Under this definition, the predicted response variable may be off by a constant intercept and a scaling factor compared to the observed values but still not suffer any loss in r^2. Whenever a model is tested on (the test set of) a library it had been trained on, it will have learned the proper intercept and scaling and so the squared Pearson coefficient r^2 is identical to the “classic” definition of R^2 in regression analysis (fraction of explained variance):

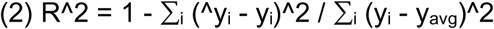

When testing on held-out libraries, however, the model has not been able to learn the bias intercept nor the scaling factor of the library’s proximal isoform logodds distribution, which will lead to overly pessimistic R^2 tests when using Equation 2 (even total covariance between predictions and observations can lead to highly negative R^2 values if predictions are off by a constant). In that case, the squared Pearson coefficient r^2 is more suitable for testing correlation.

### K-mer Linear Logistic regression

As a baseline comparison to the neural network model, we performed linear logistic regression on the combined multi-library dataset. The output response variable was the Proximal isoform ratio P^v^_Prox_ of the library variants. The input features consisted of 6-mer occurrence counts in the USE, CSE and DSE regions, and a One-hot-encoding vector of each sequence variant’s origin library.

Specifically, the input feature matrix X was constructed as follows: For each variant, the USE, CSE and DSE regions were separately scanned with a 1-nt stride for 6-mer sequence motifs. For each 6-mer, the corresponding occurrence count in the input matrix X was incremented by one. Finally, a binary library indicator feature was set to one (and encoded in matrix X) to indicate the current variant’s library origin.

The Logistic Regression model consists of a weight vector w = (w_1_, …, w_m_) of length m and a scalar bias term w0, where m is the input feature dimensionality (the total number of 6-mer features and library indicators). The predicted proximal usage ratio ^PvProx for variant v is computed as the sigmoidal activation of the weighted sum of inputs:

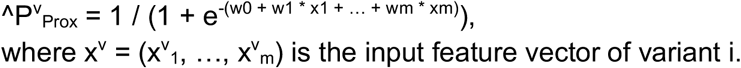

The regression weights corresponding to the library indicator features effectively encode individual per-library bias terms that absorb the library-specific mean proximal isoform logodds. By switching bias terms depending on library allows the model to capture context-specific characteristics (e.g. positioning of randomized regions, knockdown of otherwise present motifs, etc.) in those weights while keeping the 6-mer feature weights as unbiased as possible.

The model is trained by minimizing the Negative Log-Likelihood L of the library data set:

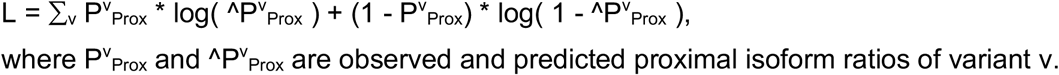

Training is done using an LM-BFGS optimization procedure from the python package SciPy (Jones et al. 2016). The model was L_2_-regularized (imposed with a loss-term ⅄ *|**w**|^2^) and cross-validated to find a suitable value for ⅄, however the optimal parameter value found was 0.

### Isoform-based DNN

The isoform-based DNN model (APARENT when trained to predict isoform ratios) is a convolutional neural network (CNN) that regresses the scalar-valued proximal isoform ratios of the library variants given a one-hot-encoding of the 186-bp long variant sequence as input. Identical to the linear logistic regression model, a one-hot-encoding of library indicator features (representing library background sequence) was also used as input.

The input data set to the CNN was constructed by one-hot-encoding the variant sequences. Specifically, for each variant sequence v of length 186, an input matrix X^v^ of size 4×186 was created, where exactly one element x^v^_k,j_ of each column j was set to one, indicating whether the nucleotide at position j is A (k = 0), C (k = 1), G (k = 2), or T (k = 3).

Architecturally, the CNN was constructed from two convolutional layers using ReLU activation functions. The first layer has 70 filters of width 8×4 and the second layer has 110 filters of width 6×1. A Max-pooling layer of grid-size 2 separates the convolutional layers. The flattened convolutional output is fed to a fully connected layer of 80 hidden units with 0.2 dropout, which finally connects to a logistic regression node that outputs the predicted proximal isoform ratio ^^^P^v^_Prox_. To account for non-sequence-related cross-library variation, a vector of bias weights indexed by the source library was added to the final regression layer. The network was trained by SGD (mini-batch stochastic gradient descent) to minimize the logloss of observed proximal isoform ratio (identical to the loss function used for linear logistic regression) using the python library Theano (The Theano Development Team et al. 2016).

Spatially close competing CSEs (occurring by chance in the randomized sequence and inserted in the USE or DSE of the proximal PAS) might impose a non-linear effect on the up- or downregulation of the primary proximal CSE, but since the neural network input covers this part of the sequence space, it will be able to learn such relationships. Far-away distal sites (not covered in the input to the network), however, are assumed to act upon APA selection independently from the proximal site and adds only an additive bias towards site preference. Given this assumption, any library-specific bias due to the composition and distance of the (non-random) distal site will be absorbed by the linear library-specific bias weights (which are indexed by the library indicator features).

### Cleavage-based DNN

In contrast to the isoform-based DNN which was trained on the scalar-valued proximal isoform ratios, the cleavage-based DNN was trained to predict the distribution of cleavage across the sequence. Specifically, the target output is the vector P^v^ of normalized cleavage counts. At the top of the network architecture is a 187-dimensional multinomial Softmax node that outputs the predicted cleavage distribution ^P^v^ = (^p^v^_1_, …, ^p^v^_186_, ^p^v^_dist_).

Architecturally, the two DNN versions are almost identical. The obvious difference is the output layer. Another difference is how library biases are handled. For the isoform-based model, a scalar bias weight was used to capture the mean isoform ratio log odds of each library. For the Cleavage-based model, a 187-dimensional vector of weights was added to the final regression layer to capture the library-specific cleavage bias at each position.

The network was trained by minimizing the cross-entropy (or multi-class logloss) between predicted and observed cleavage distributions of each variant. Because raw cut data was missing (or removed due to potential internal priming artifacts) at certain positions depending on library, each specific library was used to train the model only on a subset of softmax probability outputs. In particular, for Alien2 the model could not be trained on the cut positions at all due to the lack in knowledge of exact cut positions. Instead, for Alien2 the cleavage-model was trained on the proximal isoform ratio logloss (the isoform-based objective). The total loss L is defined as:

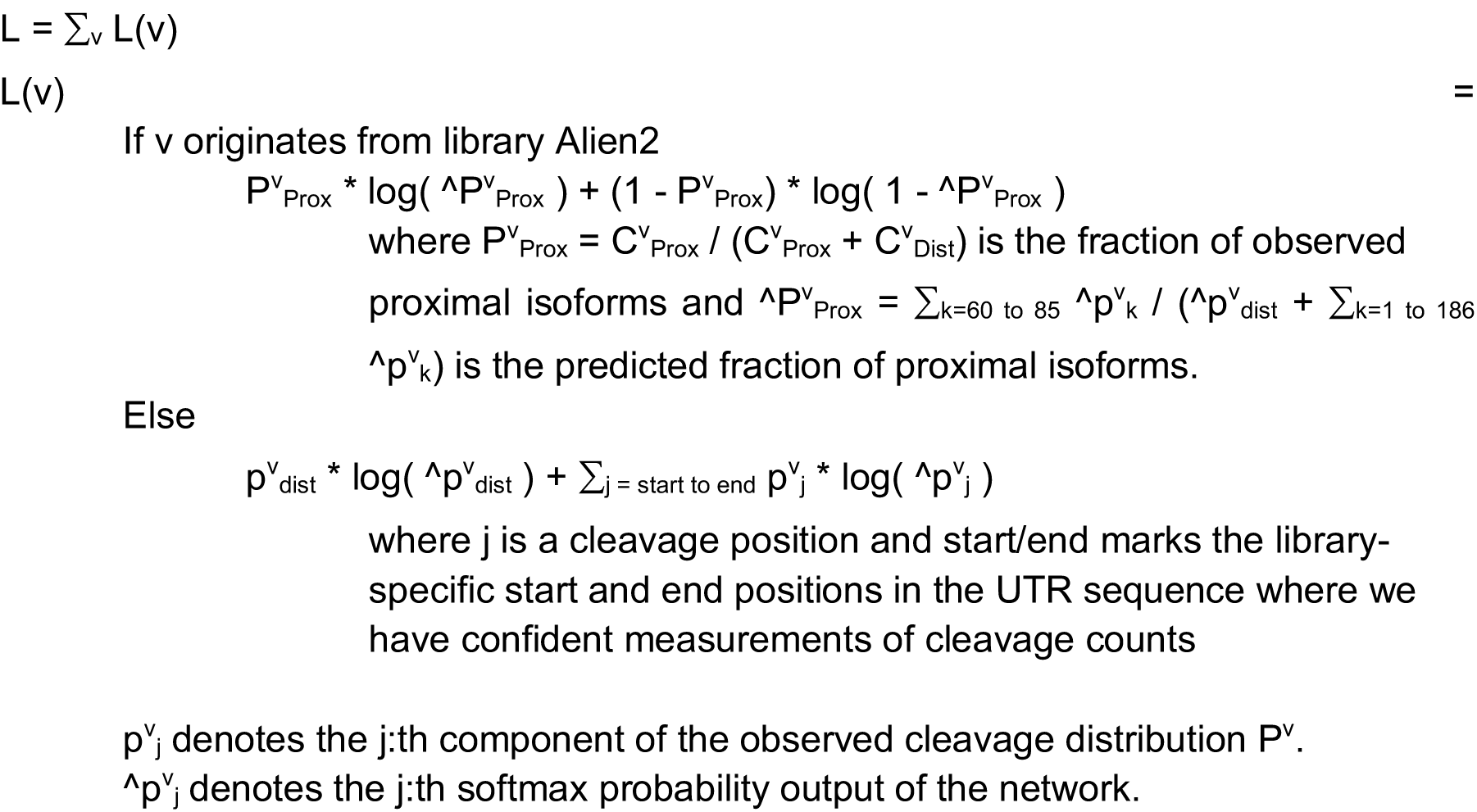

### Visualization and effect quantification of convolutional filters

The positional effect of each convolutional filter in the first layer was measured by computing correlation coefficients between filter activations at every given position and the predicted proximal isoform logodds scores (Figure S3A). The test set sequence variants from the Alien2 library were fed through the network, predicting the proximal isoform logodds logit(P^i^_Prox_) for each variant i. Next, the filter activations A^i^_k,j_ of the k:th filter at every position j were recorded for each variant i. The Pearson correlation coefficient was then calculated between filter activations (A^1^_k, j_, …, A^n^_k,j_) at position j and isoform logodds predictions (logit(P^1^_Prox_), …, logit(P^n^_Prox_)) across the test set. This coefficient value was used to color the corresponding position in the filter heatmap.

To generate consensus sequence logos for the convolutional filters representing their maximal activation, the 5,000 input subsequences of the test set that resulted in maximal filter activation were stacked into a positional weight matrix (PWM) and used to generate a sequence logo for each filter (Alipanahi et al. 2015). By purposefully biasing the selection of sequences to be maximally activating examples, the consensus logos effectively visualize the canonical form of the motif learned by a filter. To visualize the mean sequence element that a filter is sensitive to, the selection of sequences used to generate the PWM was chosen as 40,000 randomly sampled sequences from the UTR library, and their weight contribution to the PWM was scaled by the filter response.

The same framework was used to decompose the second convolutional layer into a set of sequence motifs with associated position effects (Figure S3C). After feeding the Alien2 UTR library test set through the network, layer 2 filter activations were recorded at every position and correlated with the proximal isoform logodds predictions. The subsequences of the test set resulting in maximal layer 2 filter activation were stacked into a PWM, generating a maximal-activation sequence logo for each filter.

A similar visualization scheme was used to interpret the regulatory impact on cleavage site selection of every convolutional filter in the first layer of the cleavage-based model (Figure S4A). The main difference is that, since the output predictions are multinomial cleavage site probabilities, the resulting heatmaps are two-dimensional. Using a random sample of 120,000 sequences from the Alien1 library, each filter’s activation at every position was recorded and correlated with the magnitude of cleavage at every other position, resulting in a two-dimensional heatmap. Maximal-activation sequence logos were generated for each filter as previously described.

### Visualization of higher-level network layers through optimization

Interpreting the higher-level layers of neural networks is a challenging and still open problem. One of the most commonly employed strategies in image recognition is to use gradient-based methods for optimizing an input image with respect to different components (neuron activations) of the network, thus generating visual representations that the network has learned to be sensitive towards (Simonyan et al. 2013). Consider the activation *A*^*L*^_*j*_(*I*) of a neuron *j* within network layer *L* when receiving input pattern *I*. The objective is to maximize *A*^*L*^_*j*_(*I*) by optimizing *I*. I.e., the goal is to find:

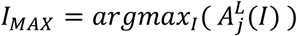

*I* is directly optimized by keeping its elements as free weights and applying backpropagation of error through the CNN (Simonyan et al. 2013). However, in that context *I* is an unconstrained image with pixel values in range (-inf, +inf). This is not the case for a sequence-based CNN; *I* is a one-hot-encoding and can only be decoded into a valid sequence if every column sums to 1 and is zero in every column component except for one. The optimization problem is hard under these constraints, however it can be simplified by approximating *I* as a column-wise softmax-activation over a matrix of free weights *W*:

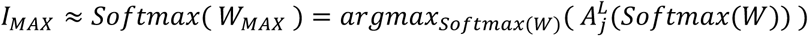

This extra Softmax-layer can easily be injected between the free weight matrix W and the first convolutional layer of the CNN (Figure S3E), which constrains the column sums to one while keeping the network differentiable w.r.t. *W*. After optimization by backpropagation through the CNN, softmax(*W*) can be interpreted as a PWM encoding of a sequence that maximally activates a particular neuron in the network. By randomly initializing W, different local maximas (and corresponding PWMs) can be found. Also, there is no need to regularize W, since the softmax transfer function removes any trace of large weights before being fed to the CNN.

The optimization routine for the fully-connected layer (illustrated in Figure S3F) was performed on 50 random sequences encoded with the Alien2 wildtype template UTR and optimizing them with respect to maximal activation of a particular neuron. The optimized sequence PWMs were then stacked together into a sequence logo.

The final proximal isoform class probability was optimized for using the same framework (Figure S3H). Starting from 50 random sequences encoded with a particular library wildtype UTR, the sequences were optimized for proximal isoform probability (the output neuron of the network). The routine was replicated for each library UTR background. Timelapses of the optimization were generated by saving the current state of the sequence PWMs at each iteration of backpropagation.

### APADB data processing and evaluation

The latest version of the human APADB dataset (Homo sapiens v2) was downloaded and 200bp up-and downstream of the cut site was extracted from the hg19 reference genome for each APA event, thus obtaining the sequence regions containing the complete PASs. The dataset comes pre-processed with read counts for each annotated isoform, so the relative isoform ratio of an APA site A versus its downstream competing site B was estimated as C_A_ / (C_A_ + C_B_) (where C_A_ and C_B_ are the mapped read count supporting each isoform). The data was filtered to retain a subset of high-quality examples; events with a total read count less than 2,000 were removed, and events with CSEs differing by more than 2 bases from the canonical hexamer AATAAA were also removed. The retained set of 674 events had their PAS sequences aligned such that the CSE hexamer was located at position 50 as expected by the model.

The isoform-based DNN was used to predict the isoform logodds of both the proximal and distal PAS sequence of each APA event. These two scalar-valued scores were used together with the natural log of the site distance to linearly regress the isoform logodds estimated from the APADB read counts (the log of the odds of C_A_ / (C_A_ + C_B_)). No regularization was used.

The extended model was evaluated with leave-one-out cross-validation, where one APA event was left out of the training data and all other events were used for training. This procedure was repeated for all of the 674 events. Each trained model was then tasked with predicting the isoform logodds of the held-out event, resulting in a test of 674 data points (Figure 5B).

An identical network and a simpler 6-mer regression model was trained exclusively on the APADB dataset and used for comparison by testing each model’s accuracy on the task of classifying an event as either mainly proximally or distally polyadenylated. For this test, the read count filter on the APADB dataset was lowered to 1,500, resulting in a set of 1,040 events. 75% (780 events) were randomly chosen as training data and the remaining 25% (260 events) were used as test data. The reason for lowering the read count filter is that more data points were needed to form an independent test set of sufficient size, as we could not perform leave-one-out cross-validation when training a full convolutional neural network (as is done for the APADB- only trained network). The ADADB-only network was trained until it was predicting the isoform logodds with zero error. Interestingly, this did not lead to overfitting (Figure S5B), as the test set error never started to increase again.

### GEUVADIS data processing

The RNA-Seq .bam-files from the GEUVADIS accession E-GEUV-1, the genotype variant call files of the 1000 Genomes project (vol 1, phase 1, v3, accession 20101123), and the Human Tandem 3’ UTR .gff annotation from the MISO project were downloaded. The Tandem 3’ UTR annotation was used to identify coordinate regions within 50 bp of annotated polyadenylation sites (cut sites) in the genome. The 1000 Genomes .vcf-files were filtered for variants occurring within these regions. The reference and variant sequences of the filtered set of variants were extracted from the hg19 reference genome. Finally, the RNA-Seq data from GEUVADIS were processed using MISO, which estimates the reference and variant isoform ratios. The set of variants were further filtered such that the isoform ratios had a narrow confidence interval (at most 25% difference between upper- and lower confidence bounds). The number of reference samples of each variant that passed the confidence filter had to be greater than 10.

This final set of variants was passed to the isoform-based model in order to predict the change in isoform logodds between the reference and variant sequence. The predicted change in isoform logodds was directly compared to the change in isoform logodds estimated by MISO (the change in the log of the odds of the estimated isoform ratios).

### ClinVar data processing

The ClinVar full release for the month 2017/09 (“ClinVarFullRelease_2017-09.xml”) was downloaded together with a historical file containing molecular consequence annotations for older variants (“molecular_consequences.txt”). A dictionary of possible consequences was built by scanning through both files (e.g. “UTR-3”, “frameshift”, “Stop-Loss”, “missense”, etc.).

The consequence dictionary was used to filter the ClinVar dataset such that only non-coding 3’ UTR variants were retained. All other variants, such as coding variants or frameshifts, were removed (pathogenicity of such variants cannot possibly be predicted by an APA model). The dataset was further filtered for variants occurring within 75 bp of an annotated cut site according to APADB. The reference UTR sequences were extracted from the hg19 reference genome and variant sequences were constructed from the coordinate annotations and variant calls in the ClinVar data. Both SNVs and InDels were extracted. The reference and variant sequences were aligned such that the CSE of the PAS was located at position 50. These sequences, together with the annotated clinical significances (benign, pathogenic or undetermined), were used to make the variant predictions of Figure 6B-C. The effect of each variant was scored using the isoform-based model. The model predicted the isoform logodds of the variant and reference sequences, and the mutation score was calculated as the difference in logodds value.

